# Hypoxia causes pancreatic β-cell dysfunction by activating a transcriptional repressor BHLHE40

**DOI:** 10.1101/2022.06.28.498031

**Authors:** Tomonori Tsuyama, Yoshifumi Sato, Tatsuya Yoshizawa, Takaaki Matsuoka, Kazuya Yamagata

**Affiliations:** Center for Metabolic Regulation of Healthy Aging (CMHA), Faculty of Life Sciences, Kumamoto University, Kumamoto 860-8556, Japan; Department of Medical Biochemistry, Faculty of Life Sciences, Kumamoto University, Kumamoto 860-8556, Japan; First Department of Internal Medicine, Wakayama Medical University, Wakayama 641-8509, Japan

## Abstract

Hypoxia can occur in pancreatic β-cells in type 2 diabetes. Although hypoxia exerts deleterious effects on β-cell function, the associated mechanisms are largely unknown. Here, we show that the transcriptional repressor basic helix-loop-helix family member e40 (BHLHE40) is highly induced in hypoxic mouse and human β-cells and suppresses insulin secretion. Conversely, BHLHE40 deficiency in hypoxic MIN6 cells or in the β-cells of *ob/ob* mice reversed the insulin secretion. Mechanistically, BHLHE40 represses expression of *Mafa*, which encodes the transcription factor musculoaponeurotic fibrosarcoma oncogene family A (MAFA), by attenuating binding of pancreas/duodenum homeobox protein 1 (PDX1) to its enhancer region. Impaired insulin secretion in hypoxic β-cells was recovered by MAFA expression. Collectively, this work identifies BHLHE40 as a key hypoxia-induced transcriptional repressor in β-cells and its implication in the β-cell dysfunction in type 2 diabetes.

## INTRODUCTION

Glucose metabolism is regulated by crosstalk between pancreatic β-cells and insulin-sensitive tissues. In case of insulin resistance, β-cells increase insulin secretion to maintain normal glucose tolerance. However, when β-cells are incapable of this task, plasma concentrations of glucose increase. Prolonged exposure to hyperglycaemia has deleterious effects on β-cell function through various mechanisms, including oxidative stress, endoplasmic reticulum (ER) stress, and inflammation and contributes to the development and progression of type 2 diabetes (1–3). Because β-cells are highly dependent on oxidative phosphorylation for adenosine triphosphate (ATP) production and insulin secretion, high glucose conditions generate intracellular hypoxia due to large amounts of oxygen consumption. Importantly, hypoxia was shown to occur *in vivo* in islets in animal models of type 2 diabetes (4–7). Like oxidative and ER stress, hypoxia leads to β-cell dysfunction and loss of β-cells, supporting the idea that hypoxia is another mechanism leading to β-cell failure in type 2 diabetes (8–11).

Hypoxia-inducible factor (HIF), a heterodimeric transcription factor consisting of an oxygen-sensitive HIF-α subunit and a constitutively expressed HIF-1β subunit, plays critical roles in the cellular responses to hypoxia (12, 13). HIF induces the expression of a number of genes necessary for adaptation to hypoxia, including those involved in glycolysis, erythropoiesis, and angiogenesis. However, hyperactivation of HIF in β-cells impairs insulin secretion by switching glucose metabolism from aerobic oxidative phosphorylation to anaerobic glycolysis (14–16), suggesting that activation of HIF underlies β-cell dysfunction and glucose intolerance in hypoxia. Besides gene induction, transcriptional repression of genes also occurs in response to hypoxia and was found to involve several transcription repressors, such as RE1 silencing transcription factor (REST), BTB and CNC homology 1 (BACH1), zinc finger E-box binding homeobox 1 (ZEB1), and inhibitor of DNA binding 2 (ID2) (17). Previously, we reported that hypoxia causes the downregulation of a number of β-cell genes involved in insulin secretion in mouse islets and MIN6 β-cells (9). However, the mechanisms of hypoxia-induced transcriptional repression in β-cells and the contribution of gene repression to β-cell dysfunction are largely unknown.

In the present study, we identified the transcriptional repressor basic helix-loop-helix family member e40 (BHLHE40) as being highly induced in β-cells under hypoxic conditions and found that it inhibits glucose-stimulated insulin secretion by suppressing transcription of *Mafa*, which encodes musculoaponeurotic fibrosarcoma oncogene family A (MAFA), a transcription factor that plays critical roles in insulin secretion. We also showed that BHLHE40 deficiency reversed decreased insulin secretion by hypoxic β-cells *in vitro* and *in vivo*. Our findings present a new scenario in which hypoxia impairs β-cell function through activation of the transcriptional repressor BHLHE40.

## RESULTS

### Global gene expression in hypoxic β-cells and islets

To assess the impact of hypoxia on global gene expression of β-cells, we first performed RNA sequencing (RNA-seq) on both mouse and human islets cultured under normal and low oxygen conditions. Approximately 5% of expressed mRNAs were significantly downregulated at least 1.5-fold in hypoxic compared with non-hypoxic islets (mouse, 20% vs 5%, respectively; human, 20% vs 2%, respectively; Supplemental Figure 1A). Consistent with our previous findings (9), under hypoxic conditions the expression of a number of β-cell genes with important roles in insulin secretion was decreased in islets (Figure 1A). Gene set enrichment analysis (GSEA) revealed that two hallmark gene sets (pancreas beta cells and oxidative phosphorylation) were significantly downregulated in both mouse and human islets in hypoxia and one hallmark gene set (hypoxia) was significantly upregulated (Figure 1B and Supplemental Figure 1B). We hypothesized that transcriptional repressors are involved in the suppression of β-cell genes under hypoxic conditions. To identify the hypoxia-sensitive repressors in β-cells, we compared the RNA-seq–based data of hypoxia-induced genes in mouse islets, human islets, and MIN6 cells and found that 25 genes were elevated (Figure 1C). By analyzing the gene ontology of these genes, we discovered that activating transcription factor 3 (ATF3) and BHLHE40 are associated with transcriptional repression (Figure 1C). In addition, BHLHE41, REST, BACH1, ID1, ID2, ZEB1/ZEB2, and SNAI1 were reported elsewhere to function as hypoxia-induced transcriptional repressors (17). Among all these repressor genes, in our study, *Bhlhe40* mRNA was the most significantly increased in hypoxic mouse islets, human islets, and MIN6 cells (Figures 1, D-F). BHLHE40 (also referred to as DEC1/SHARP2/STRA13) is a member of the basic helix-loop-helix (bHLH) family and functions primarily as a transcriptional repressor by binding to DNA at class B E-box motifs (18). It plays pivotal roles in many biological processes, including cellular differentiation, cell growth, growth arrest, circadian rhythm, immunological response, and hypoxia, but its biological function in β-cells is unknown. Accordingly, we focused on the role of BHLHE40 in hypoxic β-cells.

**Figure 1.**
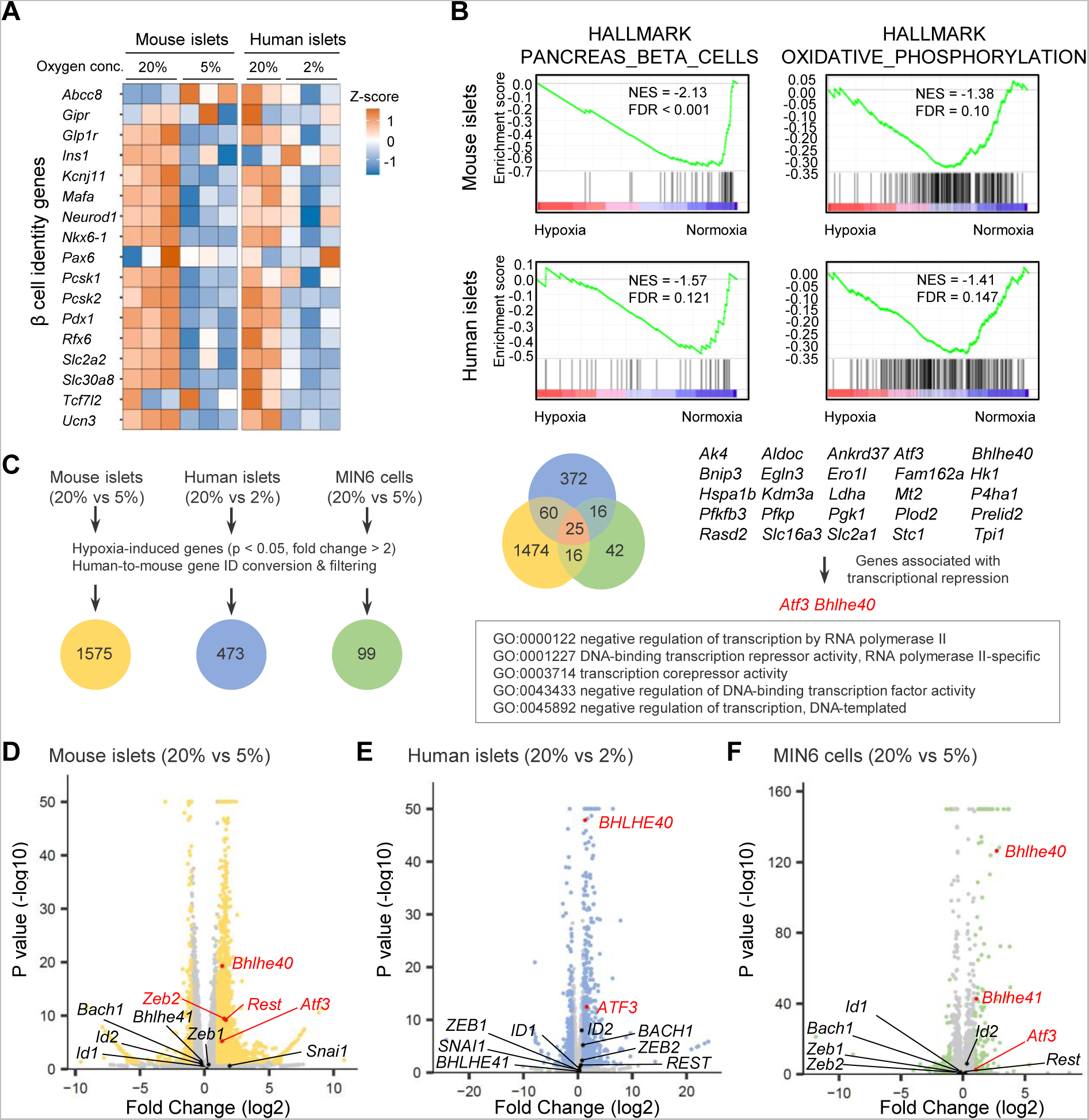
Global gene expression in hypoxic β-cells and islets. (**A**) Heatmap of β-cell genes in mouse islets (20% vs 5% O_2_ for 24 hours; n = 3) and human islets (20% [n = 2] or 2% [n = 3] O_2_ for 24 hours). (**B**) Gene set enrichment analysis of mouse islets (20% vs 5% O_2_ for 24 hours; n = 3) and human islets (20% [n = 2] or 2% [n = 3] O_2_ for 24 hours). (**C**) RNA-seq data of hypoxia-induced genes in mouse islets (20% vs 5% O_2_ for 24 hours; n = 3), human islets (20% [n = 2] vs 2% [n = 3] O_2_ for 24 hours), and MIN6 cells (20% vs 5% O_2_ for 6 hours; n = 3). The Venn diagram shows the coordinated elevation of 25 genes, two of which (*Atf3* and *Bhlhe40*) are associated with transcriptional repression. (**D**-**F**) Volcano plots showing RNA-seq data in mouse islets (20% vs 5% O_2_ for 24 hours; n = 3; **D**), human islets (20% [n = 2] vs 2% [n = 3] O_2_ for 24 hours; **E**), and MIN6 cells (20% vs 5% O_2_ for 6 hours; n = 3; **F**). *Atf3*, *Bhlhe40*, and other reported hypoxia-inducible transcriptional repressor genes (17) are shown (red, significantly upregulated genes).

### Regulation of *Bhlhe40* expression in hypoxic β-cells and islets

BHLHE40 was expressed ubiquitously in adult mouse tissues, including pancreatic islets and MIN6 cells (Figure 2A and Supplemental Figure 2A). Hypoxia rapidly increased the expression of *Bhlhe40* mRNA, i.e., within 3 hours, but a marked upregulation of BHLHE40 protein was noted at 12 hours and this upregulation persisted at 24 hours in MIN6 cells (Figure 2B and Supplemental Figure 2B). Increased BHLHE40 expression in hypoxia was also detected in mouse islets (Figure 2C and Supplemental Figure 2C).

**Figure 2.**
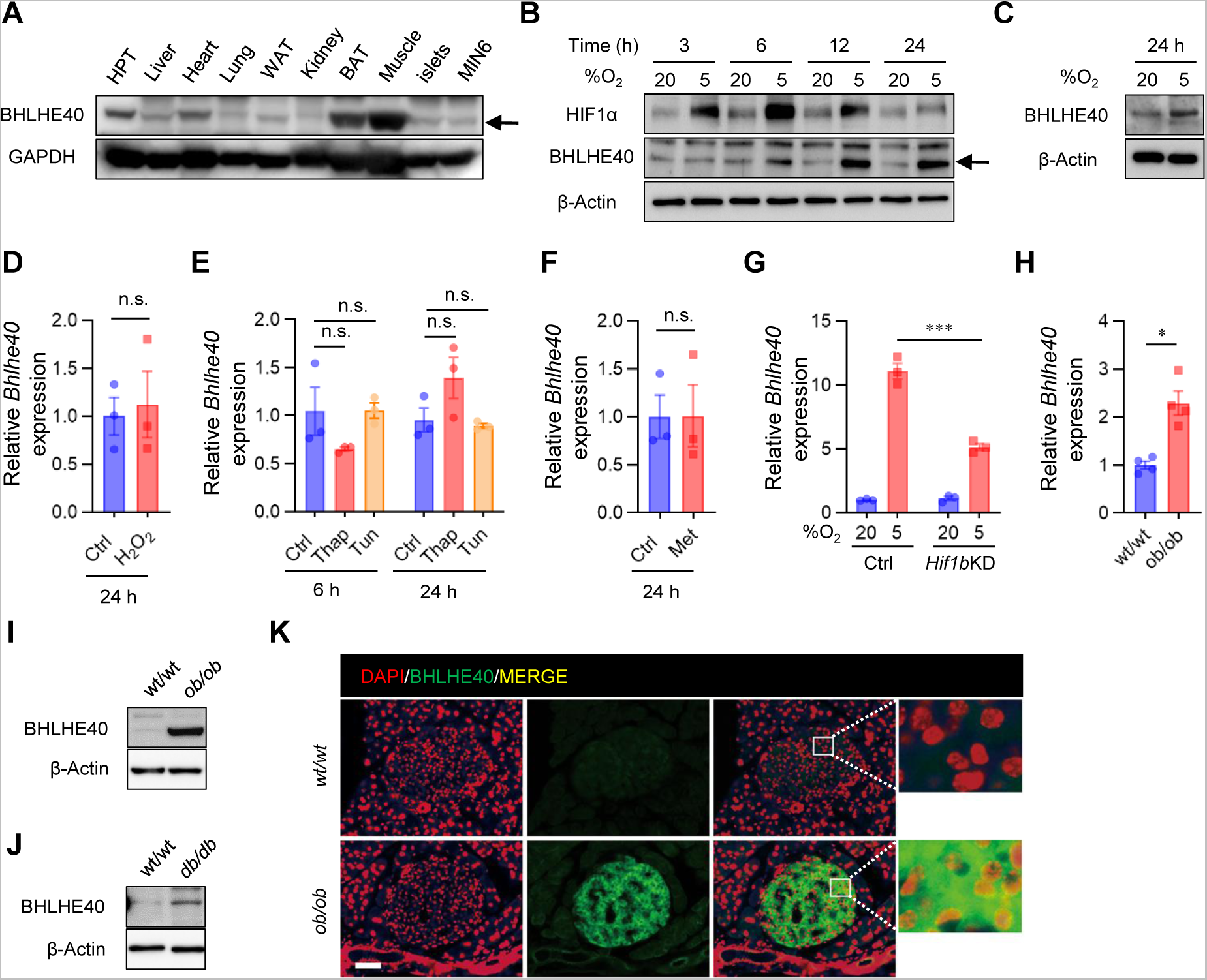
BHLHE40 expression and its regulation in hypoxic β-cells and islets. **(A-C)** Western blots of BHLHE40 expression across tissues (**A**), BHLHE40 expression in MIN6 cells cultured under 20% or 5% O_2_ for the indicated time (**B**), and BHLHE40 expression in mouse islets cultured under 20% or 5% O_2_ for 24 hours (**C**). (**D**-**F**) The effect of oxidative stress, endoplasmic reticulum stress, and energy stress on BHLHE40 expression. qRT-PCR analysis of *Bhlhe40* in MIN6 cells incubated with 10μM H_2_O_2_ (n = 3; **D**), 2μM thapsigargin (Thap) or 5 μg/ml tunicamycin (Tun) (n = 3; **E**), or 2mM metformin (Met) for 24 hours (n = 3; **F**). (**G**) The effect of short-hairpin RNA-mediated *Hif1β* knockdown (KD) on BHLHE40 expression in MIN6 cells cultured under 20% or 5% O_2_ for 24 hours (n = 3). (**H**-**K**) BHLHE40 expression in islets from diabetic mice. BHLHE40 expression was analyzed in *ob/ob* mouse islets by qRT-PCR (n = 4; **H**) and Western blotting (**I**) and in *db/db* mouse islets by Western blotting (**J**). Subcellular localization of BHLHE40 in *ob/ob* mice islets by immunohistochemical analysis (**K**). Data are mean ± SEM; *p < 0.05 and ***p < 0.001 by unpaired two-tailed Student’s *t* test. Glyceraldehyde-3-phosphate dehydrogenase (GAPDH) or β-actin was used as a loading control. Scale bar, 10 μm. HPT, hypothalamus; Ctrl, control; n.s., not significant.

Hypoxia induces oxidative and ER stress and activation of AMP-activated protein kinase (AMPK) (6, 19, 20). However, in our study, oxidative stress (caused by treatment with H_2_O_2_), ER stress (caused by treatment with thapsigargin or tunicamycin), and AMPK activation (caused by treatment with metformin) did not increase expression of *Bhlhe40* mRNA in MIN6 cells (Figures 2, D-F), suggesting that these processes are not involved in the induction of *Bhlhe40*. Previous studies showed that HIF-1 is involved in hypoxia-induced *Bhlhe40* expression (21, 22). Consistent with this finding, hypoxia-induced *Bhlhe40* mRNA expression was partially inhibited by suppression of HIF-1β (Figure 2G and Supplemental Figure 2D), indicating that hypoxia-induced *Bhlhe40* expression is partially HIF dependent in β-cells. We and others demonstrated that hypoxia occurs in islets in animal models of type 2 diabetes (4–6). Consistent with this finding, levels of *Bhlhe40* mRNA and BHLHE40 were significantly elevated (2.3-fold and 4.5-fold, respectively) in islets of *ob/ob* mice (Figures 2, H and I). Upregulation of BHLHE40 was detected also in islets of *db/db* mice (Figure 2J and Supplemental Figure 2E). Intracellular localization of BHLHE40 varies by cells (23). Immunohistochemical analysis revealed strong BHLHE40 immunoreactivity in the cytoplasm of islets in *ob*/*ob* mice, but BHLHE40 staining was clearly detected also in the nucleus of islets (Figure 2K). The nuclear localization of BHLHE40 supports its function as a transcriptional repressor.

### BHLHE40 controls insulin secretion in β-cells

To explore the role of BHLHE40 in β-cells, we generated *Bhlhe40* knockdown (KD) MIN6 cells (Supplemental Figure 3A). Although BHLHE40 is reported to be involved in cell cycle and apoptosis regulation (24), hypoxia-induced growth inhibition (Supplemental Figure 3B) and cell death (Supplemental Figure 3C) were not restored in *Bhlhe40* KD MIN6 cells. Next, we investigated the impact of *Bhlhe40* KD on glucose-stimulated insulin secretion. As described previously (9), insulin secretion by high glucose was significantly decreased under hypoxic conditions without affecting insulin content (Figures 3, A and B). Of note, *Bhlhe40* KD significantly restored hypoxia-related decreased insulin secretion (Figure 3A), and conversely, BHLHE40 overexpression significantly attenuated insulin secretion (Figure 3C and Supplemental Figure 3D). These results indicate that BHLHE40 is involved in suppressing the glucose-stimulated insulin secretion. Glucose-stimulated insulin secretion occurs after the generation of ATP through the metabolism of glucose. The increase of the ATP/ADP ratio leads to closure of ATP-sensitive K^+^ channels, membrane depolarization, an increase of cytosolic [Ca^2+^]_i_ via activation of voltage-dependent Ca^2+^ channels, and eventually the exocytosis of insulin-containing secretory granules (25).

**Figure 3.**
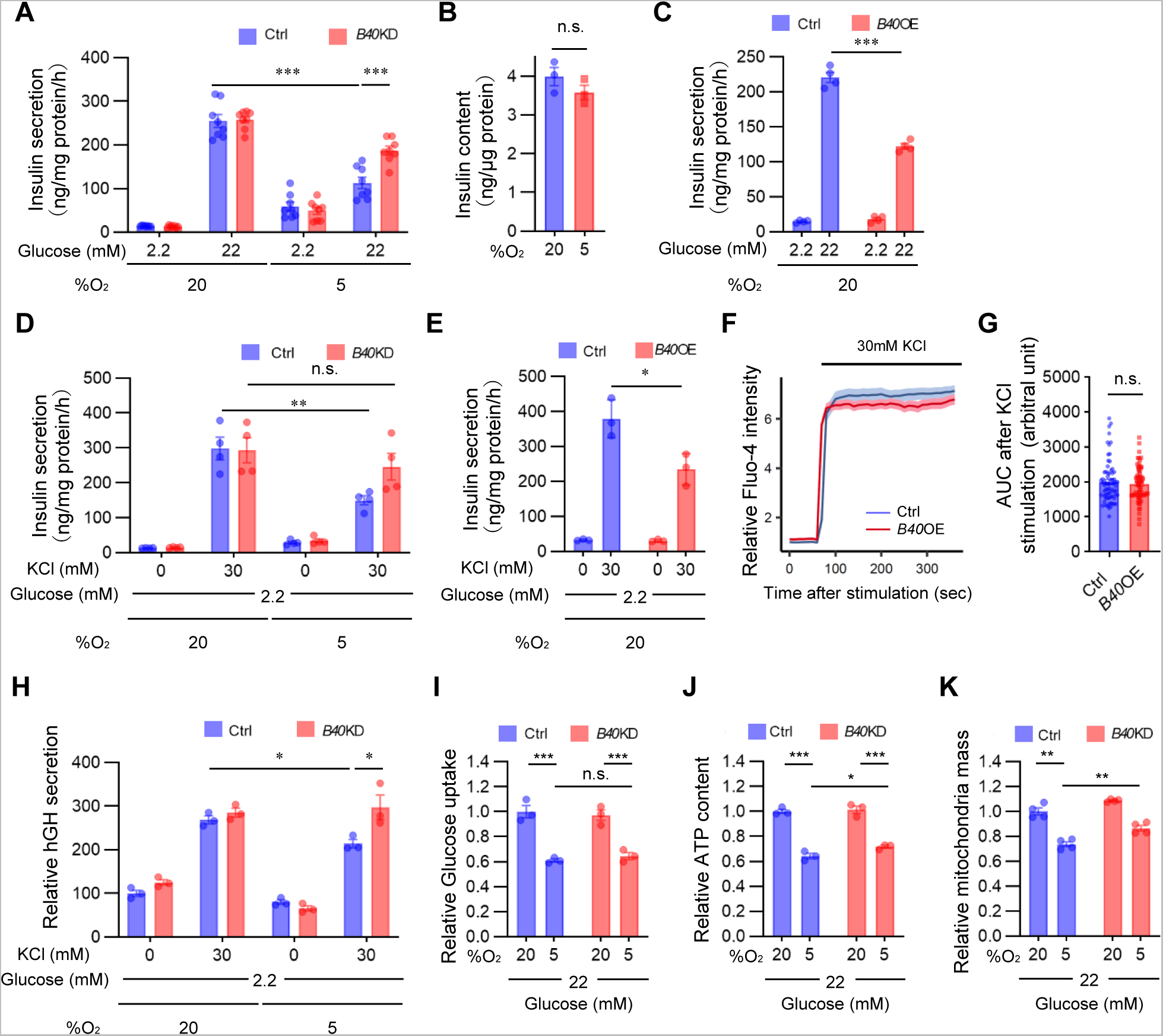
BHLHE40 controls insulin secretion in β-cells. (**A**) Glucose-stimulated insulin secretion in MIN6 cells expressing short hairpin RNA against a non-targeting Ctrl or *Bhlhe40* knockdown (*B40* KD) were cultured under 20% or 5% O_2_ for 24 hours (n = 8). (**B**) Insulin contents in MIN6 cells cultured under 20% or 5% O_2_ for 24 hours (n = 3). (**C**) Glucose-stimulated insulin secretion in MIN6 cells infected with retroviruses generated with pMx-Ctrl (Ctrl) or pMx-*Bhlhe40* (*B40* OE; n = 4). (**D**) KCl-stimulated insulin secretion in Ctrl and *B40* KD MIN6 cells cultured under 20% or 5% O_2_ for 24 hours (n = 4). (**E**) KCl-stimulated insulin secretion in Ctrl and *B40* OE MIN6 cells (n = 3). (**F**-**G**) Calcium influx stimulated by 30mM KCl in Ctrl and *B40* OE MIN6 cells (n = 70 cells from n = 3 biological replicates) (**F**) and the AUC of **F** (**G**). (**H**) hGH secretion after stimulation by 2.2mM glucose or 2.2mM glucose plus 30mM KCl in Ctrl and *B40* KD MIN6 cells cultured under 20% or 5% O_2_ for 24 hours (n = 3). (**I**-**K**) Glucose uptake (n = 3; **I**), cellular ATP content (n = 4; **J**), and mitochondrial mass (n = 4 ;**K**) in Ctrl and *B40* KD MIN6 cells cultured under 20% or 5% O_2_ for 24 hours. Data are mean ± SEM; *p < 0.05 and ***p < 0.001 by unpaired two-tailed Student’s *t* test. Ctrl, control; n.s., not significant.

High K^+^ induced membrane depolarization evokes Ca^2+^ dependent insulin exocytosis from β-cells. We next investigated the role of BHLHE40 on KCl-stimulated insulin secretion. As in the case of glucose, a significant decrease in KCl-stimulated insulin secretion was detected in hypoxic MIN6 cells but not in hypoxic *Bhlhe40* KD MIN6 cells (Figure 3D). KCl produced a significantly smaller insulin secretion in MIN6 cells overexpressing BHLHE40 than in control MIN6 cells (Figure 3E). The observation that KCl induced an equivalent increase of [Ca^2+^]_i_ levels in both cells overexpressing BHLHE40 and control cells (Figures 3, F and G) suggests that BHLHE40 affects steps after [Ca^2+^]_i_ elevation. Next, we further explored the role of BHLHE40 on exocytosis by transfecting MIN6 cells with a human growth factor (hGH) expression vector. In transfected cells, hGH is targeted to insulin-containing secretory granules, and hGH release can be used to monitor exocytosis from the cells (26, 27). As shown in Figure 3H, KCl-induced hGH secretion was significantly decreased under hypoxic conditions, and the decrease was almost completely reversed by *Bhlhe40* KD. These results emphasize the role of BHLHE40 in exocytosis in MIN6 cells.

To investigate the possibility that BHLHE40 affects multiple steps during glucose-stimulated insulin secretion, we next examined glucose uptake in *Bhlhe40* KD and control MIN6 cells. Uptake of 2-NBDG, a fluorescent derivative of glucose, was decreased by hypoxia, but *Bhlhe40* KD did not affect 2-NBDG uptake in MIN6 cells (Figure 3I), indicating that BHLHE40 does not affect glucose uptake. In line with our previous study (9), ATP levels in MIN6 cells were decreased under hypoxic conditions, and the decrease of ATP levels was significantly restored by *Bhlhe40* KD (Figure 3J). In agreement with these results, under hypoxic conditions, the decreased mitochondrial mass was significantly increased by *Bhlhe40* KD in MIN6 cells (Figure 3K). Taken together, our results show that BHLHE40 affects at least two different steps, i.e., ATP generation and exocytosis, during insulin secretion.

### BHLHE40 suppresses *Mafa* expression in β-cells

To further understand how BHLHE40 regulates insulin secretion, we performed RNA-seq analysis in control MIN6 cells and MIN6 cells overexpressing *Bhlhe40*. Compared with control MIN6 cells, MIN6 cells overexpressing *Bhlhe40* showed 2630 differentially expressed genes (1288 downregulated and 1342 upregulated; adjusted p < 0.01) (Figure 4A). Gene ontology analysis revealed that the downregulated genes included transcription genes with critical roles during insulin secretion, including *Mafa*, *Nkx2-2*, *Ppargc1a, Vdr*, *Nkx6-1*, and *Neurod1* (Figures 4, A and B). Quantitative real-time polymerase chain reaction (qRT-PCR) in independent samples confirmed the significant downregulation of *Mafa*, *Nkx2-2*, *Ppargc1a*, and *Vdr* expression by *Bhlhe40* overexpression (Figure 4C). To validate that BHLHE40 functions as a repressor of these genes, we next examined their expression in *Bhlhe40* KD MIN6 cells under hypoxic conditions. Hypoxia significantly decreased *Mafa*, *Ppargc1a, Vdr,* and *Nkx6-1* mRNA, but downregulation of *Mafa* was completely restored by KD of BHLHE40 (Figure 4D). At the protein level, suppression of MAFA by BHLHE40 and restoration of hypoxia-induced downregulation of MAFA by BHLHE40 deficiency were also detected (Figures 4, E and F). MAFA regulates genes required for insulin exocytosis, including *Stxbp1* (encoding MUNC18-1), *Napa* (encoding N-ethylmaleimide-sensitive factor attachment protein), *Syt7* (encoding synaptotagmin 7), and *Stx1a* (encoding syntaxin1A) (28–33). Under hypoxic conditions, *Bhlhe40* KD also significantly restored the downregulation of *Stxbp1*, *Napa*, *Syt7*, and *Stx1a* (Figure 4G). MAFA plays a critical role in both glucose- and KCl-stimulated insulin secretion (28, 34). Intriguingly, the glucose- and KCl-related decreased insulin secretion in hypoxic conditions was significantly restored by adeno-associated virus (AAV)-mediated overexpression of *Mafa* (Figures 4, H and I, and Supplemental Figure 4). These results indicate that BHLHE40 suppresses insulin secretion, at least in part, by reducing the expression of MAFA in β-cells.

**Figure 4.**
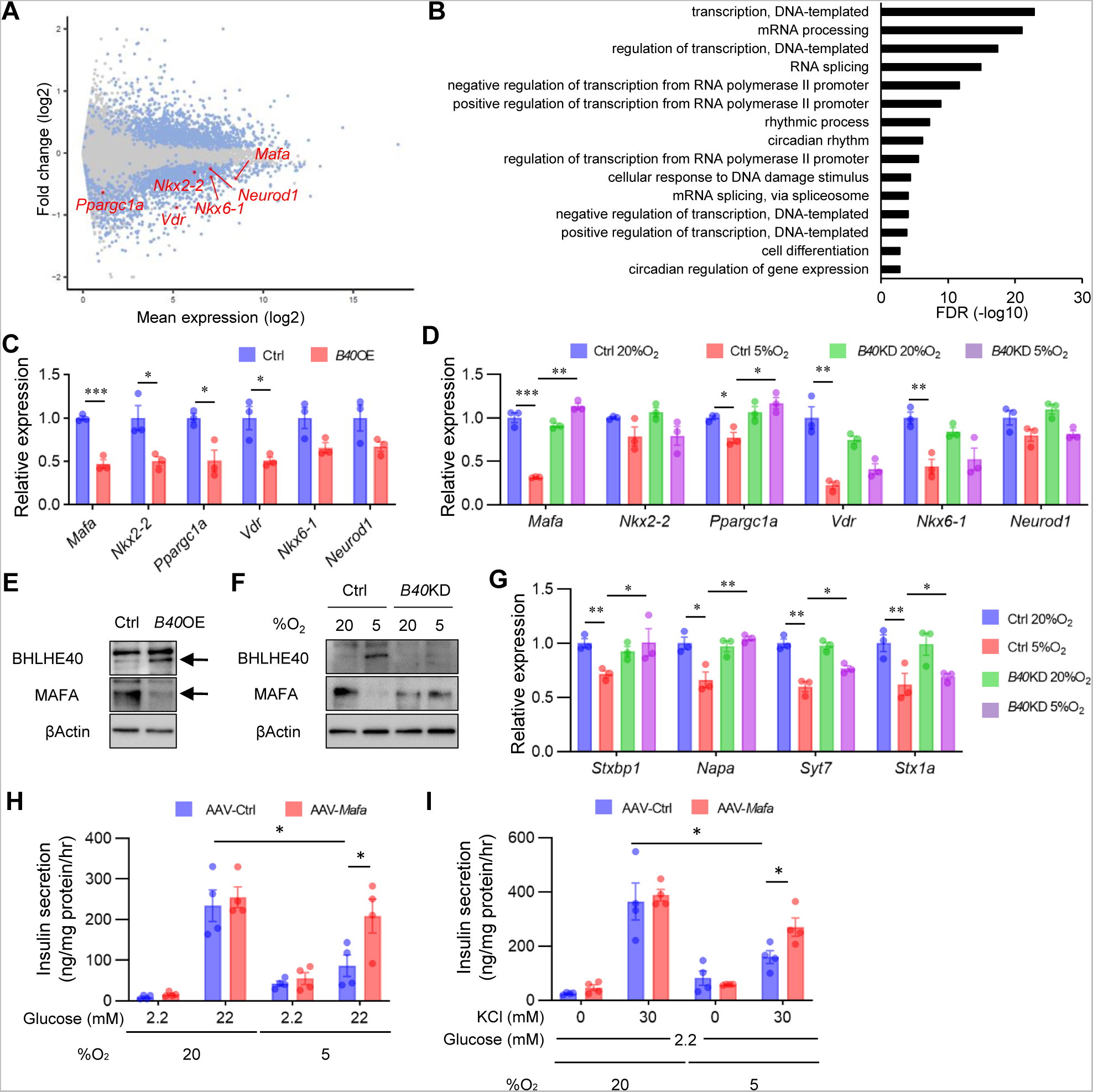
BHLHE40 suppresses *Mafa* expression in β-cells. (**A**) MA plot of RNA-seq data in Ctrl and *B40* OE MIN6 cells. Differentially expressed genes (DEGs; adjusted p value < 0.01) are shown in blue, and others in gray. DEGs functioning as β-cell transcription factors are shown in red (n = 3). (**B**) Gene ontology analysis of RNA-seq data. Downregulated DEGs were used as input. (**C** and **D**) Expression of DEGs shown in **A** was confirmed by qRT-PCR in Ctrl and *B40* OE MIN6 cells (n = 3; **C**) or Ctrl and *B40* KD MIN6 cells cultured under 20% or 5% O_2_ for 24 hours (n = 3; **D**). (**E**-**F**) Western blot of BHLHE40 and MAFA expression in Ctrl and *B40* OE MIN6 cells (**E**) or Ctrl and *B40* KD MIN6 cells cultured under 20% or 5% O_2_ for 24 hours (**F**). (**G**) qRT-PCR of MAFA target genes in Ctrl and *B40* KD MIN6 cells cultured under 20% or 5% O_2_ for 24 hours (n = 3). (**H**) Glucose-stimulated insulin secretion in MIN6 cells infected with AAV-green fluorescent protein (GFP) (Ctrl) or AAV-*Mafa* and cultured under 20% or 5% O_2_ for 24 hours (n = 4). (**I**) KCl-stimulated insulin secretion in MIN6 cells infected with AAV-GFP (Ctrl) and AAV-*Mafa* and cultured under 20% or 5% O_2_ for 24 hours (n = 4). Data are mean ± SEM; *p < 0.05, *p < 0.01 and ***p < 0.001 by unpaired two-tailed Student’s *t* test. β-actin was used as a loading control. Ctrl, control.

Peroxisome proliferator-activated receptor-γ coactivator 1-α (PGC-1α), which is encoded by *Ppargc1a*, regulates mitochondrial biogenesis and oxidative phosphorylation (35). Previous studies revealed that BHLHE40 acts as a transcriptional repressor of *Ppargc1a* (36, 37). Consistent with this finding, *Ppargc1a* expression was significantly downregulated by *Bhlhe40* overexpression in MIN6 cells, and the hypoxia-induced downregulation of *Ppargc1a* was recovered by KD of *Bhlhe40* (Figures 4, C and D), indicating that *Ppargc1a* is another target gene of BHLHE40 in MIN6 cells. In these experimental conditions, BHLHE40 KD did not affect the expression of *Nkx2-2*, *Vdr, Nkx6-1,* or *Neurod1* mRNAs (Figure 4D).

### BHLHE40 controls *Mafa* expression *via* two E-box sites in the enhancer region

To further explore the mechanism by which BHLHE40 suppresses *Mafa* expression in β-cells, we performed a reporter gene assay with a reporter plasmid. We included the mouse *Mafa* enhancer/promoter region (−10427 to +22 bp to transcriptional start site) because it shows maximum *Mafa* promoter activity in β-cells (38, 39). *Mafa* reporter gene activity was significantly decreased under hypoxic conditions, but the reduction was abolished by *Bhlhe40* KD (Figure 5A). BHLHE40 binds to E-box sequence (5’-CANNTG-3’) to suppress its target genes. Screening of the JASPAR database (40) revealed four E-box sites (A, −9909/−9899; B, −8705/−8695; C, −6987/−6976; and D, −4949/4938, relative score > 0.9) within the −10427/+22 region (Figure 5B). Overexpression of *Bhlhe40* suppressed activity of the reporter gene in MIN6 cells, but mutation of the A or C site in the reporter gene abolished the reduction of transcriptional activity by BHLHE40 (Figure 5C). Consistent with this finding, suppression of the reporter gene activity by hypoxia also was attenuated by mutation of these two sites (Figure 5D). Furthermore, chromatin immunoprecipitation (ChIP) assay revealed enhanced binding of BHLHE40 to A and C sites in MIN6 cells under hypoxic conditions (Figure 5E). These results indicate that BHLHE40 suppresses *Mafa* expression by binding to the A or C site. BHLHE40 is reported to repress transcription of the target genes by recruiting histone deacetylase (HDAC; ref. 41, 42). However, treatment with trichostatin A (TSA), an HDAC inhibitor, failed to affect the reduced reporter gene activity in hypoxic MIN6 cells (Figure 5F).

**Figure 5.**
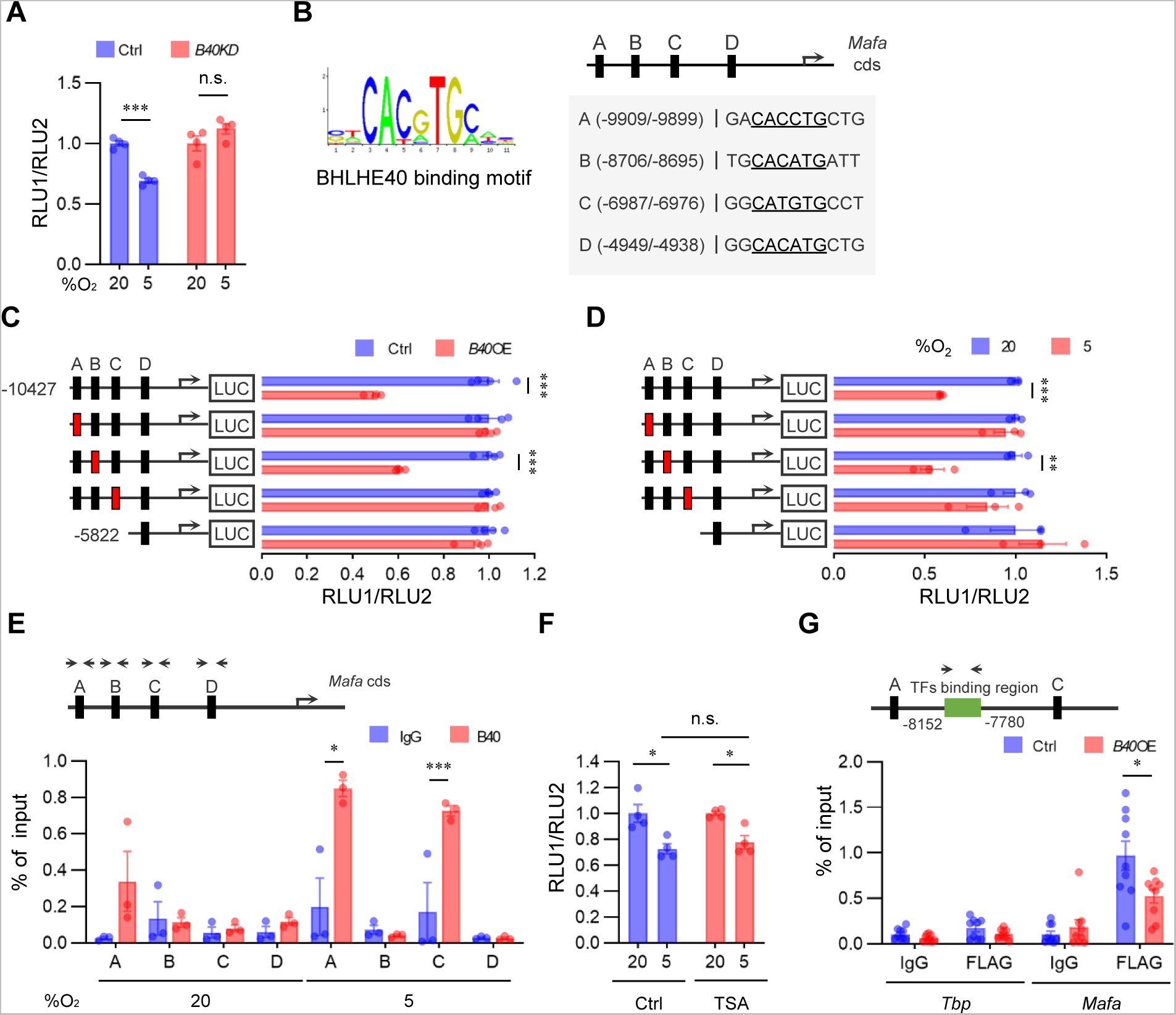
BHLHE40 controls *Mafa* expression via two E-box sites in the enhancer region. (**A**) Reporter gene analysis with luciferase plasmid infused with mouse MAFA promoter/enhancer (−10427/+22 bp from transcription start site) in Ctrl and *B40* KD MIN6 cells cultured under 20% or 5% O_2_ for 24 hours (n = 4). (**B**) BHLHE40-binding motif (upper left) and JASPAR results (lower left) are presented. JASPAR predicted four E-box sites for BHLHE40 binding on MAFA promoter/enhancer (relative score > 0.9, E-box sites are underlined). (**C**) Luciferase reporter assay was performed with MIN6 cells transfected with BHLHE40 expression plasmids, pRL-SV40 plasmid, and pGL3-Mafa plasmids (wildtype [black] and E-box mutated [red] sites; n = 4). (**D**) Luciferase reporter activity in MIN6 cells cultured under 20% or 5% O_2_ for 24 hours (n = 3). (**E**) MIN6 cells were cultured in 20% or 5% O_2_ for 24 hours, and then the proteins were immuno-precipitated by IgG or anti-BHLHE40 specific antibody, after which qRT-PCR was performed for the indicated regions (n = 3). (**F**) Luciferase reporter assay in MIN6 cells incubated with 0.1μM TSA or vehicle for 24 hours (n = 4). (**G**) Proteins sampled from Ctrl and *B40* OE MIN6 cells with FLAG-Pdx1 expression were immuno-precipitated by IgG or anti-FLAG antibody, after which qRT-PCR was performed for the indicated regions (n = 9). Data are mean ± SEM; *p < 0.05 **p < 0.01, and ***p < 0.001 by unpaired two-tailed Student’s *t* test. Ctrl, control; n.s., not significant.

The transcription factor pancreas/duodenum homeobox protein 1 (PDX1) was reported to regulate *Mafa* expression in β-cells by binding to the enhancer region (−8152 to −7780 relative to the transcription start site) (38, 43). Therefore, we investigated whether BHLHE40 represses *Mafa* expression by inhibiting PDX1 binding. Intriguingly, the ChIP assay revealed that BHLHE40 significantly reduced PDX1 binding to the *Mafa* gene in MIN6 cells (Figure 5G). These results suggest that BHLHE40 controls *Mafa* expression, at least in part, by affecting the binding of PDX1.

### Deficiency of BHLHE40 improves hyperglycaemia in *ob/ob* mice

To evaluate the role of BHLHE40 *in vivo*, we generated β-cell–specific BHLHE40 knockout (*βB40*KO) mice by crossing Pdx1-Cre mice (44) with floxed *Bhlhe40* (*Bhlhe40^fl^*) mice (Supplemental Figures 5, A and B). Body weight (Supplemental Figure 5C) and nonfasting blood glucose concentration (Supplemental Figure 5D) were similar in *βB40*KO and *Bhlhe40^fl/fl^* mice, and the intraperitoneal glucose tolerance test also showed no differences in blood glucose levels among Pdx1-Cre, *Bhlhe40^fl/fl^*, and *βB40*KO mice (Figure 6A).

**Figure 6.**
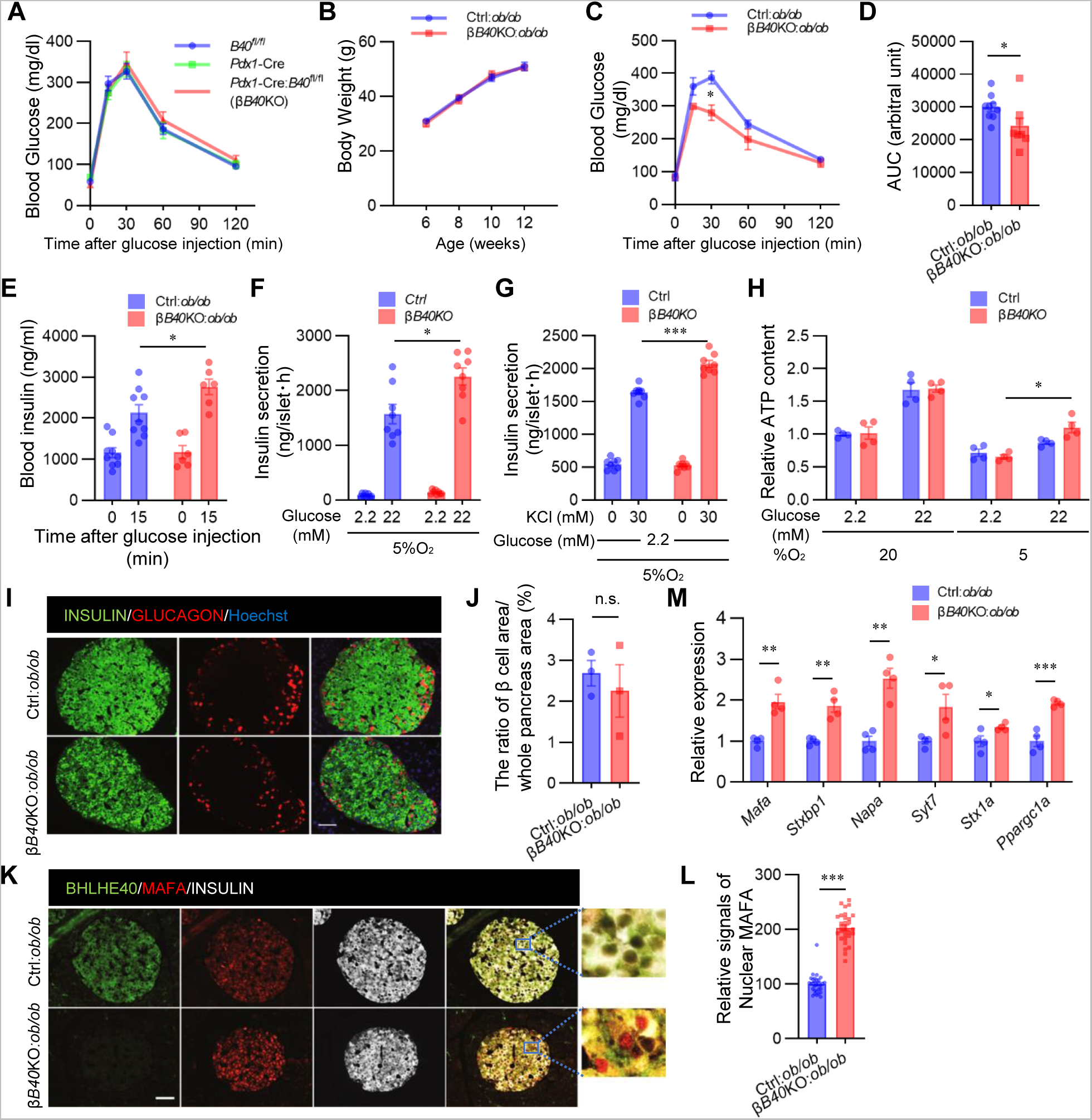
Deficiency of BHLHE40 improves hyperglycaemia in *ob/ob* mice. (**A**) Glucose tolerance test of *Pdx1*-Cre (*Cre*), *Bhlhe40^fl/fl^* (*B40^fl/fl^*), and *Pdx1*-Cre:*Bhlhe40^fl/fl^* (*Cre:B40^fl/fl^*) mice (n = 7, n = 12, and n = 9 respectively; 8-12 weeks old). (**B**) Body weight of *Bhlhe40^fl/fl^* (Ctrl):*ob/ob* and *Pdx1*-Cre:*Bhlhe40^fl/fl^* (βB40KO):*ob/ob* mice (n = 9 and n = 6, respectively). (**C** and **D**) Glucose tolerance test of Ctrl:*ob/ob* and βB40KO:*ob/ob* mice (n = 9 and n = 8, respectively; 6 weeks old) (**C**) and AUC (**D**). (**E**) Glucose-stimulated insulin secretion in Ctrl:*ob/ob* and βB40KO:*ob/ob* mice (n = 9 and n = 6, respectively; 8 weeks old). (**F**) Glucose-stimulated insulin secretion in isolated islets from Ctrl and βB40KO mice after culture under 5% O_2_ for 24 hours (n = 8). (**G**) KCl-stimulated insulin secretion in isolated islets from Ctrl and βB40KO mice after incubation with 5% O_2_ for 24 hours (n = 8). (**H**) ATP content in isolated islets of Ctrl and βB40KO mice after incubation with 20% or 5% O_2_ for 24 hours (n = 4). (**I** and **J**) Representative images of pancreatic islets stained for insulin and glucagon in Ctrl:*ob/ob* and βB40KO:*ob/ob* mice (12 weeks old) (**I**). The ratios of total islet area to whole pancreas area (%) are shown (n = 3; **J**). (**K** and **L**) Representative images of pancreatic islets stained for insulin, MAFA, and BHLHE40 in Ctrl:*ob/ob* and βB40KO:*ob/ob* mice (12 weeks old) (**K**). Fluorescence intensities of nuclear and cytosolic MAFA in **K** were quantified (n = 30; **L**). (**M**) qRT-PCR of *Mafa* and its target genes in Ctrl:*ob/ob* and βB40KO:*ob/ob* mice (n = 4). Data are mean ± SEM; *p < 0.05 **p < 0.01, and ***p < 0.001 by unpaired two-tailed Student’s *t* test. Scale bar, 10 μm. Ctrl, control; n.s., not significant.

BHLHE40 negatively regulates insulin secretion, and BHLHE40 expression was markedly upregulated in *ob/ob* pancreatic islets (Figure 2I). We then investigated the effects of β-cell–specific BHLHE40 deficiency in *ob/ob* mice (*βB40*KO:*ob/ob* mice). In these mice, BHLHE40 deficiency in β-cells had no effect on obesity (Figure 6B) or insulin sensitivity (Supplemental Figure 5E). However, the mice displayed better glucose tolerance than control *Bhlhe40^fl/fl^*:*ob*/*ob* (control:*ob*/*ob*) mice (Figures 6, C and D). In agreement with the results obtained in MIN6 cells, glucose-stimulated insulin secretion was significantly increased in *βB40*KO:*ob/ob* mice (Figure 6E). Insulin secretion by high glucose (Figure 6F) and KCl (Figure 6G) was also increased in *βB40*KO islets under hypoxic conditions. We measured the ATP content with 2.2mM and 22mM glucose under hypoxic conditions and found that ATP levels were significantly increased in *βB40*KO islets compared with control islets (Figure 6H). There was no significant difference in the ratio of β-cell area to whole pancreas area between *βB40*KO:*ob/ob* and control:*ob*/*ob* mice (Figures 6, I and J), but stronger nuclear immunostaining of MAFA was detected in *βB40*KO:*ob/ob* mice (Figures 6, K and L). Lastly, the increased expression of *Mafa, Stxbp1*, *Napa*, *Syt7*, *Stx1a*, and *Ppargc1a* in *βB40*KO:*ob/ob* islets was confirmed by qRT-PCR (Figure 6M). Taken together, the results show that BHLHE40 deficiency improves glucose tolerance in *ob/ob* mice by enhancing insulin secretion.

## DISCUSSION

Adaptation to hypoxia involves 3 major responses: increased glycolysis to cope with ATP depletion, increased oxygen delivery, and inhibition of energy-demanding processes such as gene transcription (17). HIF transcriptional factors are known to play central roles in glycolytic ATP production and oxygen delivery, but the mechanisms underlying transcriptional repression in hypoxia are poorly understood. By screening hypoxia-induced genes in mouse and human islets and MIN6 cells, we showed that the transcriptional repressor BHLHE40 is highly induced in hypoxic β-cells. We also demonstrated that BHLHE40 negatively regulates insulin secretion by suppressing transcription of *Mafa*. Hypoxia is involved in β-cell dysfunction, and the contribution of HIF-1 to this process is well established (7, 14–16). However, our present findings present a novel scenario in which hypoxia decreases insulin secretion by inducing BHLHE40 (Figure 7).

**Figure 7.**
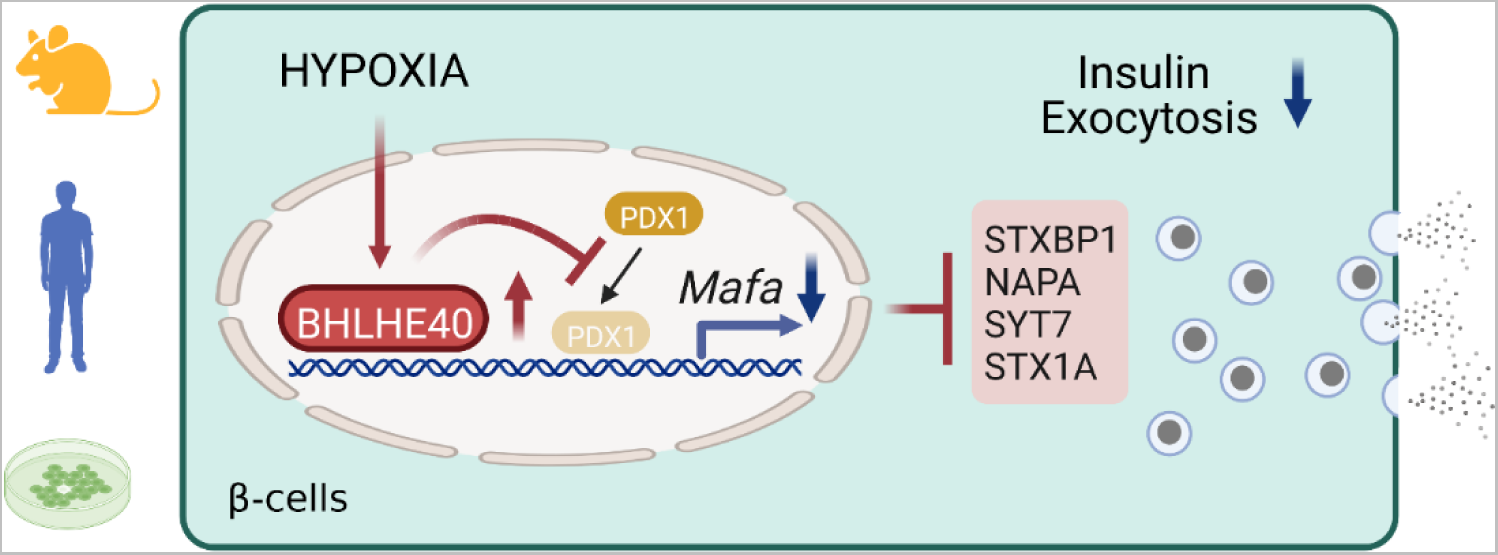
A proposed model for how hypoxia causes β-cells dysfunction.

BHLHE40 was previously reported to suppress transcription of target genes by recruiting HDACs (37, 45). In contrast, we found that TSA treatment did not relieve the BHLHE40-mediated repression of *Mafa*, indicating that *Mafa* repression by BHLHE40 is independent of the recruitment of HDAC. On the other hand, we demonstrated that BHLHE40 inhibits the binding of PDX1 to the critical enhancer region of *Mafa*. Repressors are reported to regulate transcription by interacting with activator proteins (46). BHLHE40 might suppress *Mafa* transcription by interacting with PDX1. Alternatively, BHLHE40 might change DNA conformations by recruiting chromatin-remodeling factors. The family of large Maf proteins comprises MAFA, MAFB, and MAF (c-Maf). Intriguingly, recent studies demonstrated that BHLHE40 represses the expression of *Mafb* and *Maf* mRNAs in macrophages (47). Thus, BHLHE40 seems to be a common repressor of the large-Maf family. Further studies are necessary to clarify the mechanism by which BHLHE40 suppresses these genes. Moreover, we also revealed that BHLHE40 suppresses transcription of *Ppargc1a* and decreases ATP levels in β-cells. Because PGC-1α plays important roles in ATP production (48, 49), it is plausible that the decreased expression of PGC-1α may be involved in the reduced ATP levels and impaired insulin secretion under hypoxic conditions.

In the present study, we showed that approximately 5% of genes were downregulated in hypoxic islets. In addition to BHLHE40, we also found an increased expression of ATF3 in β-cells in hypoxia. Previously, ATF3 was reported to function as a transcriptional repressor and to suppress genes related to glucose metabolism, such as *Irs2*, *Nrf1*, and *Pparg* (50, 51). Thus, hypoxia might affect β-cell function through not only BHLHE40 but also other transcriptional repressors, such as ATF3.

In conclusion, we identified BHLHE40 as a novel hypoxia-induced transcriptional repressor that negatively regulates insulin secretion in β-cells. Because β-cell dysfunction in type 2 diabetes is progressive, new approaches to slow the progression are needed. Inhibition of BHLHE40 might be a new therapeutic strategy for preventing the β-cell dysfunction by hypoxia.

## METHODS

### Mouse models

C57B6 wild-type (WT), *ob/ob*, and *db/db* mice were purchased from KBT Oriental Co., Ltd. (Saga, Japan). Mice carrying the Bhlhe40*^tm1a(KOMP)Wtsi^*allele (C57BL/6NTac-Bhlhe40tm1a(KOMP)Wtsi/WtsiPh, EM:09819) were obtained from the European Conditional Mouse Mutagenesis Program (EUCOMM) and crossed with *FLPe* mice (B6-Tg(CAG-FLPe)36, RBRC01834, RIKEN BRC, Ibaraki, Japan) to remove a LacZ reporter and a Neo cassette flanked by two Frt sites. To achieve a *Bhlhe40* deletion in β-cells, Bhlhe40*^fl/fl^*mice were further crossed with Pdx1-Cre mice (gift from Dr. Douglas A. Melton). β-cell–specific Bhlhe40 knockout mice on *ob/ob* background were generated by crossing with Bhlhe40*^fl/fl^*:*ob/+* mice and Pdx1-Cre:Bhlhe40*^fl/fl^*:*ob/+* mice. Mice were housed under a 12-hour light/dark cycle with free access to water and normal chow (CE-2; CLEA, Tokyo, Japan). Room temperature was maintained at 22 ± 1-2°C.

### Human pancreatic islets

Human islets were commercially purchased from Prodo Laboratories (Irvine, CA). The cadaveric donor had no history of diabetes (32-year-old male; BMI, 25.1; HbA1c, 5.1%). Islets were cultured according to the Prodo Laboratories instructions.

### Cell lines

MIN6 cells were gifts from Jun-ichi Miyazaki (Osaka University). They were maintained in Dulbecco’s modified Eagle’s medium (DMEM) supplemented with 10% (v/v) fetal bovine serum (FBS), 0.1% (v/v) penicillin/streptomycin (P/S), and 50μM β-mercaptoethanol at 37℃ in 5% CO_2_, 95% air. For the hypoxic cell culture, a multi-gas incubator (APM-300; ASTEC, Fukuoka, Japan) was used. Retroviral packaging cell line Platinum-E (Plat-E; RV-101) cells were purchased from Cell Biolabs, Inc. (San Diego, CA). They were maintained in DMEM supplemented with 10% (v/v) FBS and 0.1% (v/v) P/S at 37℃ in 5% CO_2_, 95% air. 293AAV cells were purchased from Cell Biolabs Inc. (AAV-100). They were maintained in DMEM with sodium pyruvate supplemented with 10% (v/v) FBS, 1x Glutamax, 1x Opti-MEM, and 0.1% (v/v) P/S at 37℃ in 5% CO_2_, 95% air.

### Plasmids

The HA-tagged mouse *Bhlhe40* coding sequence was excised from a pCAGGS-DEC1 plasmid (RDB08473, Riken, Saitama, Japan; ref. 52) and subcloned into pcDNA3.1 and pMXs-Puro Retroviral vector (RTV-012, Cell Biolabs, Inc.). For knockdown experiments, oligonucleotide encoding Bhlhe40 short hairpin RNA (shRNA; target sequence: 5’-GCACGTGAAAGCATTGACA-3’; ref. 53) and Hif1β shRNA (target sequence: 5’-GGACAGAGATCCAAGGTTT-3’) were cloned into the pSIREN-RetroQ expression vector (631526; Clontech Laboratories Inc., Mountain View, CA). The FLAG-tagged mouse *Pdx1* coding sequence was amplified by PCR and subcloned into a pMXs-Neo Retroviral vector (RTV-011, Cell Biolabs, Inc.). pGL3-basic-MafA plasmid (−10427/+22 from transcription start site) was previously reported (39). The pGL3-basic-MafA plasmids with single deletion of the E-box site at −9910/−9899 (A; GAAAAATGCTG), −8706/−8695 (B; TGAAAATGATT), or −6987/−6976 (C; GGAAAATGCCT) were generated with a KOD-Plus Mutagenesis Kit (SMK-101, TOYOBO, Osaka, Japan). The truncated pGL3-basic-MafA plasmid (−5811/+22) was generated by BglII digestion and re-ligation. The mouse *Mafa* coding sequence was amplified by PCR and subcloned into pAAV-MCS (VPK-410, Cell Biolabs. Inc.).

### MIN6 cells stably overexpressing/silencing a target gene

To generate stable overexpression cell lines, retroviral vectors (pMx-Puro-HA-*Bhlhe40*, pMx-Puro Control, pMx-Neo FLAG-*Pdx1* or pMx-Neo Control plasmid) were transfected into Plat-E cells with JetPRIME transfection reagent (114-15, Polyplus, New York, NY), and MIN6 cells were infected with the respective retroviruses and selected by incubation with puromycin (5 μg/ml) for 2 days or G418 (500 μg/ml) for 4 weeks. For stable knockdown of *Bhlhe40*, pSIREN-RetroQ-Bhlhe40 or pSIREN-RetroQ-control vector was transfected into Plat-E cells, and MIN6 cells were infected with the retroviruses and selected by incubation with puromycin (5 μg/ml) for 2 days.

### AAV-*Mafa* preparation

293AAV cells were plated onto 10-cm dishes. At 80% confluency, cells were transfected with 2 μg/dish pAAV-DJ, 3 μg/dish pHelper, and either 2 μg/dish pAAV-GFP or pAAV-*Mafa* with JetPRIME transfection reagent. Three days after transfection, AAV-GFP and AAV-*Mafa* were purified with the AAV pro Extraction Solution Kit (6235, Takara, Shiga, Japan), and the titers were determined with a Quick Titer AAV Quantification Kit (VPK-145, Cell Biolabs, Inc.) according to the manufacturer’s instructions.

### Isolation of mouse islets

Mice were euthanized by cervical dislocation and subjected to bile duct cannulation and digestion of the pancreas with a mixture of collagenase P (11-249-002-001, Roche, Basel, Switzerland), hyaluronidase (H3506; Sigma-Aldrich, St. Louis, MO), and protease inhibitor cocktail (Nacalai Tesque, Kyoto, Japan) for 25 to 30 minutes in a warm (37℃) water bath. Isolated islets were collected manually. Islets were maintained in RPMI-1640 supplemented with 10% (v/v) FBS, 0.1% (v/v) P/S, 50μM β-mercaptoethanol, 10mM HEPES, and 1mM sodium pyruvate at 37℃ in 5% CO_2_, 95% air.

### qRT-PCR

MIN6 cells were homogenized in Sepasol-RNA I reagent (09379-55, Nacalai Tesque), and RNA was manually isolated by phenol-chloroform extraction and ethanol precipitation. RNA from isolated islets was prepared with the RNeasy Micro Kit (74004, QIAGEN, Hilden, Germany) according to the manufacturer’s instructions. cDNA was synthesized with a Prime Script RT Reagent Kit (RR047A, Takara Bio Inc., Shiga, Japan). qRT-PCR was performed with SYBR Premix Ex TaqII (RR820A, Takara Bio Inc.) in an ABI 7300 thermal cycler (Applied Biosystems, Foster City, CA). All data were normalized to *Actb* or *Tbp*. The primers for this study are listed in Supplemental Table 1.

### RNA-seq analysis

RNA was extracted with the RNeasy Micro Kit (74004, QIAGEN) according to the manufacturer’s instructions. For Figure 1 samples, sequencing libraries were prepared with a NEBNext Ultra II Directional RNA Library Prep Kit (7765L, New England Biolabs, Ipswich, MA), and samples were sequenced on an Illumina NextSeq 500 platform in 76bp single-end reads. For Figure 4 samples, sequencing libraries were prepared with a NEBNext Ultra II RNA Library Prep Kit (E7770, New England Biolabs) and samples were sequenced on an Illumina NovaSeq 6000 platform in 150bp paired-end reads. For reanalysis of RNA-seq data of *db/db* mice islets (accession number: GSE 107489), raw RNA-seq data were downloaded from NCBI Sequence Read Archive and converted to the fastq format with SRA-Tools (v2.10.9). Reads were trimmed for universal Illumina adaptors with TrimGalore (v0.6.5) (http://www.bioinfor5matics.babraham.ac.uk/projects/trim galore/) and then mapped to GENCODE 36 genome sequence (for human) or M25 genome sequence (for mouse) with HISAT2 (v2.2.1; ref. 54). Mapped reads were sorted and converted to a binary alignment/map format with SAMtools (v1.11; ref. 55). Gene assembly and quantification were performed with Stringtie (v2.1.4; ref. 56), and gene-level count matrixes were generated with python script prepDE.py3 (http://ccb.jhu.edu/software/stringtie/dl/prepDE.py3). Differentially expressed genes (DEGs) were determined with DESeq2 (v1.28.0; ref. 57). DEGs (adjusted p-value < 0.01) were used for gene ontology analysis with David (v6.8; ref. 58, 59). Raw and processed RNA sequencing were deposited in the Gene Expression Omnibus (GEO) under accession number GSE202603.

### Screening for hypoxia-induced genes associated with transcriptional repression

Hypoxia-induced genes were defined as genes with an adjusted p value of less than 0.05 and a fold change (hypoxia/normoxia) greater than 2 in DESeq2 outputs. To identify hypoxia-induced genes that are commonly listed in mouse and human islets and MIN6 cells and are related to transcriptional repression, DESeq2 outputs in each group were processed as follows: For human islets, human-to-mouse Ensembl gene identifiers (IDs) conversion was carried out with Ensembl BioMart; and for MIN6 cells and mouse islets, only genes with mouse Ensembl gene IDs that can be mapped to those in human were selected. After this gene IDs conversion and gene filtering, overlapped genes were determined. Genes associated with transcriptional repression were manually selected out of the overlapped genes based on gene ontology annotations against each gene (obtained from Ensembl BioMart).

### GSEA

Gene set enrichment analysis was performed with GSEA (v4.03; ref. 60) for “pre-ranked” analyses, with the fold change between normoxia samples and hypoxia samples as the input. Mouse Ensembl IDs were converted to be compatible with the human annotations of the MSigDB gene lists by using a Mouse ENSEMBL Gene ID to Human Orthologs MSigDB.v7.4.chip.

### Western blotting

Cells were lysed in RIPA buffer (50mM Tris-HCl [pH 8.0], 150mM NaCl, 0.1% sodium dodecyl sulfate [SDS], 1% NP-40, 5mM EDTA, and 0.5% sodium deoxycholate) with a protease inhibitor cocktail. Total proteins were separated by SDS polyacrylamide gel electrophoresis, transferred to polyvinylidene difluoride membranes (Immuno-P; Millipore, Bedford, MA) and then probed with the primary antibody. After incubation with the horseradish peroxidase (HRP)-conjugated secondary antibodies, the HRP signals were visualized by using Chemi-Lumi One Super (02230-30, Nacalai Tesque) and a ChemiDocTM Imaging System (Bio-Rad Laboratories, Hercules, CA). The primary antibodies used in this study were anti-β-actin antibody (M177-3, MBL), anti-BHLHB2 antibody (H00008553-M01, Abnova), anti-MafA antibody (A300-611A, Bethyl Laboratories, Montgomery, TX), anti-glyceraldehyde-3-phosphate dehydrogenase (GAPDH) antibody (2118, Cell Signaling Technology, Danvers, MA), anti-HIF1α antibody (NB100-479, Novus Biologicals, Centennial, CO), and anti-HIF1β antibody (5537, Cell Signaling Technology).

### Insulin secretion assay and insulin content in MIN6 cells

MIN6 cells were seeded in a 24-well plate. In Figures 3A, 3B, and 3D, cells were incubated in 20% or 5% O_2_ for 24 hours before assay. Cells were pre-conditioned in low-glucose (2.2mM) Krebs-Ringer-bicarbonate HEPES (KRBH) buffer (120mM NaCl, 4.7mM KCl, 1.2mM KH_2_PO_4_, 2.4mM CaCl_2_, 1.2mM MgCl_2_, 20mM NaHCO_3_, 10mM HEPES, and 0.5% (v/v) BSA) for 1 hour. Cells were washed once with low-glucose KRBH and incubated in low-glucose KRBH for 1 hour, and the supernatant was collected. Then, cells were stimulated in high-glucose (22mM) KRBH or low-glucose + KCl (30mM) KRBH for 1 hour, and the supernatant was collected. Next, cells were lysed in cell lysis buffer, and the protein concentration was measured with a Pierce BCA Protein Assay Kit (23225, Thermo Fisher Scientific, Waltham, MA) to normalize the insulin level. To measure insulin content, MIN6 cells were pelleted and resuspended in acid-ethanol (1.5% HCl in 70% EtOH), rotated overnight at 4℃ and neutralized with 1M Tris-HCl (pH 7.5; 1:1). The insulin concentration was determined with a mouse insulin enzyme-linked immunosorbent assay (ELISA; TMB) kit (AKRIN-011T; Shibayagi Co., Ltd., Gunma, Japan). A hypoxia chamber glove box (Creative Bio Station: CBS-120; ASTEC) was used to achieve continuous hypoxic conditions during the assay.

### Insulin secretion assay in mouse islets

The islets isolated from Ctrl and βB40KO mice were cultured in 5% O_2_ for 24 hours. Then, they were preincubated for 30 minutes in KRBH buffer containing 2.2mM glucose. For glucose challenge, they were incubated in KRBH buffer containing 2.2mM or 22mM glucose for 30 minutes, and the supernatant was collected; and for KCl challenge, they were incubated in KRBH buffer containing 2.2mM or 2.2mM glucose plus 30mM KCl for 30 minutes, and the supernatant was collected. The insulin concentration was determined with a mouse insulin ELISA (TMB) kit (AKRIN-011T and AKRIN-011S; Shibayagi Co., Ltd.). A hypoxia chamber glove box (Creative Bio Station: CBS-120; ASTEC) was used to achieve continuous hypoxic conditions during the assay.

### Cell proliferation assay

Before the assay, Ctrl or *Bhlhe40* KD MIN6 cells were seeded in a 96-well plate at 2 x 10^4^ cells/well, incubated overnight and transferred to 20% or 5% O_2_. Cells were counted at 0, 24, 48, and 96 hours with a Cell Counting Kit-8 (343-07623, Dojindo, Kumamoto, Japan), and the absorbance (450/655) was measured by an iMark microplate reader (Bio-Rad Laboratories).

### Cell death assay

Before the assay, Ctrl or *Bhlhe40* KD MIN6 cells were transferred to and cultured in 20% or 5% O_2_ for 24 hours. Then, they were incubated in 0.5 μg/ml PI (341-07881, Dojindo) for 10 minutes, after which flow cytometric analyses were performed with a FACSCalibur (BD Biosciences, Franklin Lakes, NJ) and FlowJo software (Tomy Digital Biology, Tokyo, Japan).

### Calcium assay

Ctrl or *Bhlhe40* OE MIN6 cells were plated on glass-bottomed culture dishes (627871, Greiner Bio-One, Frickenhausen, Germany). The cells were preincubated with KRBH buffer for 45 minutes and then incubated with KRBH buffer containing 2.2mM glucose, 2μM Fluo4-AM (F311, Dojindo), 0.02% Pluronic F-127 (P2443, Sigma-Aldrich), 2.5mM probenecid (162-26112, Wako Pure Chemical Industries, Ltd.), and Hochest 33258 (343-07961, Dojindo) for 30 minutes, after which the buffer was replaced with dye-free KRBH buffer. To perform the KCl challenge, buffer was changed by hand-aspirating it and gently adding an equal amount of KRBH buffer containing 2.2mM glucose and 30mM KCl back into the dish with a micropipette Time-series images were acquired every 10 seconds with a fluorescent microscope (BZ-X700; Keyence, Osaka, Japan) and analyzed with Keyence software.

### hGH secretion assay

Ctrl or *Bhlhe40* KD MIN6 were transfected with either pcDNA3-empty or pcDNA3-hGH. At 24 hours after transfection, cells were transferred to and cultured in 20% or 5% O_2_ for a further 24 hours. Cells were preincubated in glucose-free KRBH for 15 minutes and then incubated in KRBH with or without KCl (30mM) for 30 minutes, and the supernatant was collected. Cells were then lysed in cell lysis buffer, and the protein concentration was measured with a Pierce BCA Protein Assay Kit (23225, Thermo Fisher Scientific) to normalize the hGH level. The hGH concentration was measured with Human Growth Hormone ELISA kit (ab190811, Abcam, Cambridge, UK).

### Glucose uptake assay

Before the assay, Ctrl or *Bhlhe40* KD MIN6 cells were transferred to and incubated in 20% or 5% O_2_ for 24 hours. Cells were preincubated in KRBH for 15 minutes and then incubated in KRBH containing 200μM 2-NBDG (23002-v, Peptide Institute, Inc., Osaka, Japan) plus 22mM glucose for 15 minutes, after which flow cytometric analyses were performed with FACSCalibur (BD Biosciences) and FlowJo software (Tomy Digital Biology).

### Mitochondrial mass assay

Before the assay, Ctrl or *Bhlhe40* KD MIN6 cells were transferred to and incubated in 20% or 5% O_2_ for 24 hours. Then, cells were incubated in 2nM nonyl acridine orange (A-1372, Invitrogen, Carlsbad, CA) for 15 minutes, after which flow cytometric analyses were performed with FACSCalibur (BD Biosciences) and FlowJo software (Tomy Digital Biology).

### Luciferase assay

MIN6 cells were transiently transfected with firefly luciferase plasmid (either pGL3-basic-MafA or its derivatives) and renilla luciferase plasmid (pRL-SV40) with jetPRIME transfection reagent (114-15, Polyplus). For *Bhlhe40* overexpression experiments, Bhlhe40 expression plasmids (either pcDNA3.1-empty or pcDNA3.1-Bhlhe40) were additionally transfected. Forty-eight hours after transfection, cells were lysed and assayed with firefly luciferase and renilla luciferase substrates in the Dual-Luciferase Reporter Assay System (E1980, Promega). Firefly luciferase activity (RLU1) was normalized to renilla luciferase activity (RLU2). For hypoxia experiments, cells were transferred to 20% or 5% O_2_ for 24 hours before the luciferase activities were measured.

### ChIP assay

MIN6 cells were fixed in 1% formaldehyde for 10 minutes at room temperature, and then the reaction was quenched by 150mM glycine for 5 minutes. The fixed cells were incubated in 0.5% Nonidet P-40 lysis buffer for 15 minutes on ice, and the nuclei were pelleted and incubated in SDS lysis buffer (50mM Tris-HCl [pH 8.0], 1% SDS, 10mM EDTA). Chromatin was then sheared with a Bioruptor sonicator (C30010016, Diagenode, Seraing, Belgium) by 10 cycles of sonication at 30 seconds on, 30 seconds off. The sheared chromatin was diluted 5-fold in ChIP dilution buffer (50mM Tris-HCl [pH 8.0], 167mM NaCl, 1.1% Triton X-100, and 0.11% sodium deoxycholate) and then incubated in Dynabeads protein A (1001D, Invitrogen) and protein G (1003D, Invitrogen) for 1 hour at 4℃. After removing the beads, the chromatin was incubated in 4 μg of anti-Bhlhe40 antibody (NB100-1800, Novus Biologicals) or control IgG (2729, Cell Signaling Technology) overnight at 4℃. The antibody-protein complexes were isolated by incubation with magnetic beads (Invitrogen Dynabeads protein A and protein G) for 6 hours at 4℃. Then, samples were sequentially washed with low-salt RIPA buffer (50mM Tris-HCl [pH 8.0], 150mM NaCl, 1mM EDTA, 0.1% SDS, 1% Triton X-100, and 0.1% sodium deoxycholate), high-salt RIPA buffer (50mM Tris-HCl [pH 8.0], 500mM NaCl, 1mM EDTA, 0.1% SDS, 1% Triton X-100, and 0.1% sodium deoxycholate), LiCl wash buffer (10mM Tris-HCL [pH 8.0], 250mM LiCl, 1mM EDTA, 0.5% Nonidet P-40, and 0.5% sodium deoxycholate), and Tris-EDTA buffer and finally eluted and reversely cross-linked in ChIP direct elution buffer (50mM Tris-HCl [pH 8.0], 5mM EDTA, and 0.5% SDS) overnight at 65℃. DNA was then extracted and collected by phenol-chloroform extraction and ethanol precipitation. DNA was amplified by qRT-PCR with SYBR Premix Ex Taq II (RR820A, Takara) in ABI 7300 thermal cycler (Applied Biosystems) with the primers listed in Supplemental Table 1.

### Metabolic analysis of mice

Male 6- to 12-week-old mice were used for metabolic analysis. For the glucose tolerance test, mice were fasted overnight. After intraperitoneal glucose administration (2 g/kg for wildtype background, 1 g/kg for *ob/ob* background), blood glucose levels were measured at 0, 15, 30, 60, 90, and 120 minutes. For the insulin tolerance test, mice were fasted for 4 hours. After intraperitoneal insulin administration (1 unit/kg for wildtype background, 3 units/kg for *ob/ob* background), blood glucose levels were measured at 0, 30, 60, 90, and 120 minutes. For the glucose-stimulated insulin secretion assay, mice were fasted overnight. After intraperitoneal administration of 3 g/kg glucose, blood samples were collected at 0 and 15 minutes. The plasma insulin level was determined with a mouse insulin ELISA (TMB) kit (AKRIN-011S; Shibayagi Co., Ltd.).

### Immunohistochemistry and β-cell mass assessment

Pancreas tissues from mice were fixed with 10% (v/v) neutral buffered formalin (060-01667, Wako Pure Chemical Industries, Ltd.) for 16 to 18 hours at 4°C. Fixed samples were embedded in paraffin, cut into 4-μm cross-sections, mounted on MAS-coated slides (Matsunami Glass, Osaka, Japan). The deparaffinized sections were subjected to antigen retrieval (110℃, 20 minutes) with HistoVT One (L6F9587, Nacalai Tesque) and stained with the following primary antibodies: anti-insulin antibody (A0564, 1:400; Dako, Santa Clara, CA), anti-glucagon antibody (ab92517, 1:400; Abcam), anti-DEC1/BHLHE40 antibody (NB100-1800, 1:100; Novus Biologicals), and anti-MafA antibody (A300-611A, 1:100; Bethyl Laboratories). After reaction with fluorescent dye-conjugated secondary antibodies, fluorescent signals were captured with an all-in-one fluorescent microscope (BZ-X700; Keyence). The total islet area (μm^2^) composed of insulin- and glucagon-stained cells was measured by Keyence software, and the ratio of the total islet area to the total pancreas area was calculated.

### Statistics

The significance of differences was assessed by unpaired two-sided Student’s *t* tests, unless stated otherwise. All data are presented as means ± SEM. No statistical analysis was used to predetermine the sample size.

### Study approval

The handling and killing of mice were performed in compliance with the animal care guidelines of Kumamoto University. All animal experiments were conducted in accordance with the guidelines of the Institutional Animal Committee of Kumamoto University and were approved by the Kumamoto University Ethics Review Committee for Animal Experimentation (ID: A29-001, A 2019-048, A 2021-001). Human islets experiments were approved by the Ethical Committee of Kumamoto University Graduate School of Medical Sciences (No. 2389).

## AUTHOR CONTRIBUTIONS

T.T., Y.S. and K.Y. conceived and designed the work; T.T. and Y.S. obtained the data; T.T., Y.S., T.Y., T.M., and K.Y. analyzed the data; T.T., Y.S. and K.Y. drafted the manuscript; All authors reviewed the results and approved the final version of the manuscript.

## ACKNOWLEDGEMENT

We thank Dr. Douglas A. Melton (Harvard University) for providing Pdx1-Cre mice and Dr. Shingo Usuki (Kumamoto University) for his technical support. We also thank the members of Dr. Yamagata’s laboratory for their technical assistance and discussions. This study was supported by Grants-in-Aid for Scientific Research (B) (19H03711 and 22H03129; K.Y.), by a Grant-in-Aid for Challenging Research (Exploratory) (19K22639; K.Y.), and by a Grant-in-Aid for Scientific Research (C) (19K09008; Y.S.); by grants from the Takeda Science Foundation (K.Y. and Y.S.); and by grants from the program for Leading Graduate Schools HIGO (T.T.) and Center for Metabolic Regulation of Healthy Aging (T.T.).

## Notes

**Conflict-of-interest** The authors have declared that no conflict of interest exists.

### Competing Interest Statement

The authors have declared no competing interest.

