## Supplemental Materials for "Hypoxia causes pancreatic β-cell dysfunction by activating a transcriptional repressor BHLHE40"

**Supplemental material.**

**Supplemental Figures:**

**Supplemental Figure 1 (related to Figure 1)**

**Supplemental Figure 2 (related to Figure 2)**

**Supplemental Figure 3 (related to Figure 3)**

**Supplemental Figure 4 (related to Figure 4)**

**Supplemental Figure 5 (related to Figure 6)**

**Supplemental Table 1**

**A**

| | | Mouse islets (20% vs 5%) | Human islets (20% vs 2%) | MIN6 cells (20% vs 5%) |
|---|---|---|---|---|
| Fold change > 1.5 P < 0.05 | up | 2231/21166 (10.5%) | 1162/20019 (5.8%) | 420/16036 (2.6%) |
| | down | 1067/21166 (5.0%) | 1116/20019 (5.6%) | 358/16036 (2.2%) |

**B**

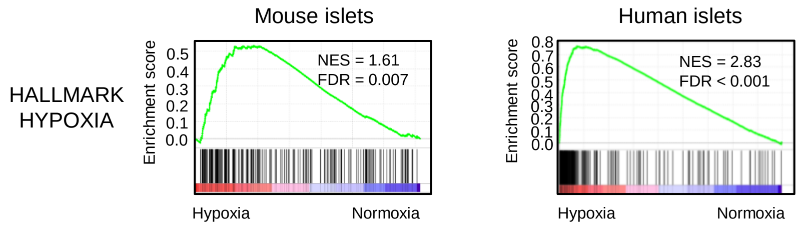


**Supplemental Figure 1 (related to Figure 1). Hypoxia caused extensive transcriptional upregulation and repression in β-cells.** (**A**) The number and the ratio of the genes that were significantly upregulated or downregulated (fold change > 1.5, $p < 0.05$) by hypoxia in mouse islets (20% vs 5% $O_2$ for 24 hours, n = 3), human islets (20% [n = 2] vs 2% [n = 3] $O_2$ for 24 hours), and MIN6 cells (20% vs 5% $O_2$ for 6 hours, n = 3). (**B**) Gene set enrichment analysis of RNA-seq data showed that the hypoxia gene set was significantly upregulated in both hypoxic mouse and human islets.

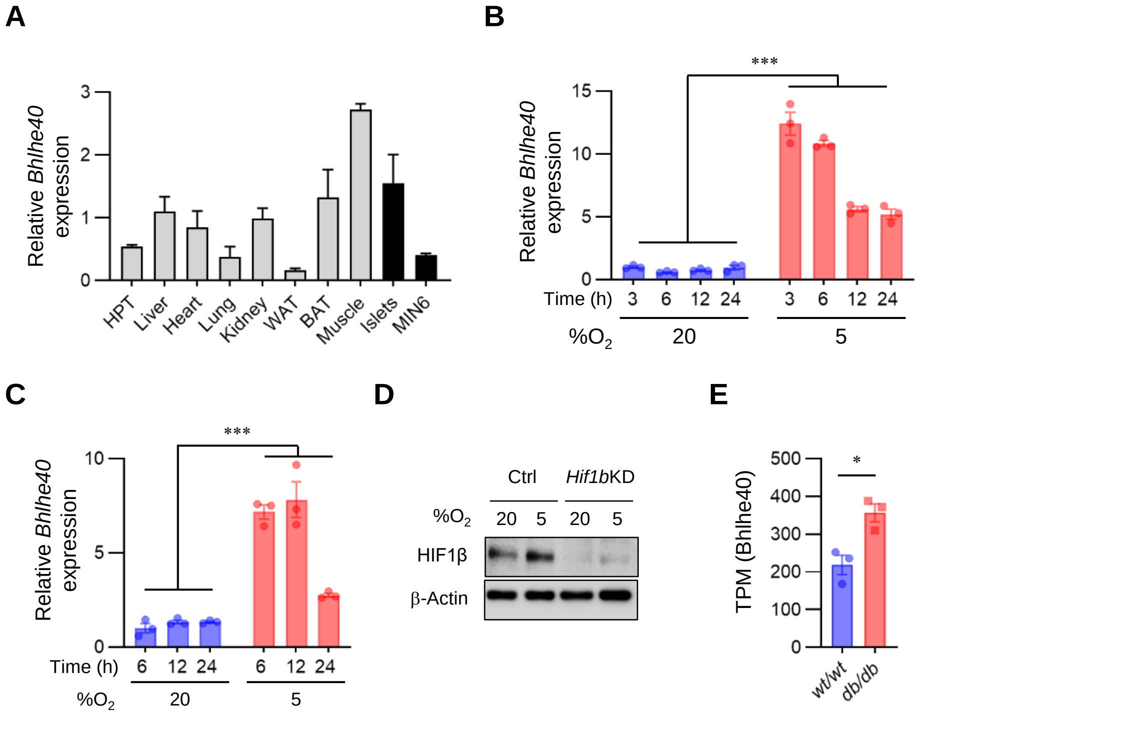


**Supplemental Figure 2 (related to Figure 2). Elevated BHLHE40 expression in hypoxic and diabetic β-cells.** (**A**) *Bhlhe40* expression across tissues by qRT-PCR (n = 3). (**B** and **C**) *Bhlhe40* expression in MIN6 cells (n = 3; **B**) and mouse islets (n = 3; **C**) cultured under 20% or 5% $O_2$ for the indicated time assessed by qRT-PCR. (**D**) MIN6 cells expressing shRNA against a non-targeting Ctrl or *Hif1β* were cultured under 20% or 5% $O_2$ for 6 hours, and then the protein samples were collected. Knockdown (KD) of *Hif1β* was confirmed by Western blotting. (**E**) The re-analysis of previously published RNA-seq data of *db/db* mouse islets showed significant upregulation of *Bhlhe40* (n = 3). Data are mean ± SEM; *p < 0.05 and ***p < 0.001 by unpaired two-tailed Student's *t* test. Ctrl, control. HPT, hypothalamus.

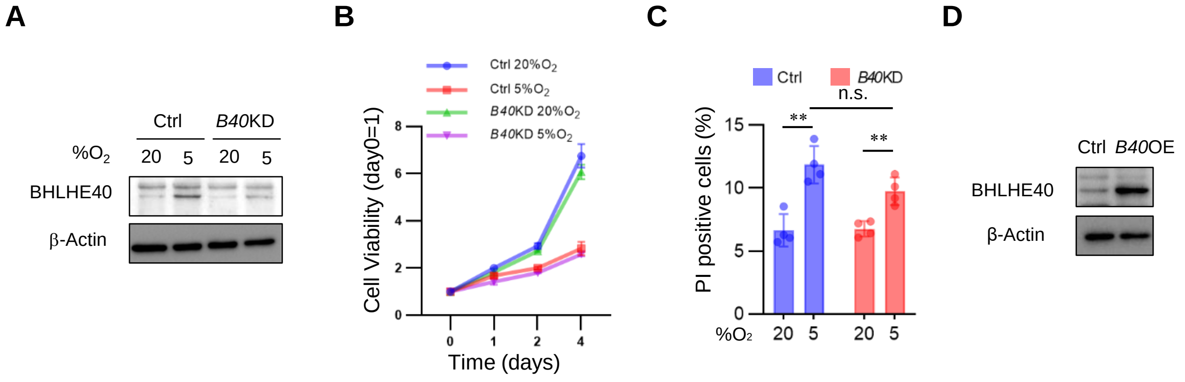


**Supplemental Figure 3 (related to Figure 3). BHLHE40 was not involved in cell proliferation or cell death caused by hypoxia.** (**A**) Confirmation of *Bhlhe40* knockdown by Western blotting in control (Ctrl) and *Bhlhe40* knockdown (*B40* KD) MIN6 cells cultured under 20% or 5% $O_2$ for 24 hours. (**B**) Cell proliferation in Ctrl and *B40* KD MIN6 cells cultured under 20% or 5% $O_2$ for the indicated times (n = 6). (**C**) Cell death in Ctrl and *B40* KD MIN6 cells cultured under 20% or 5% $O_2$ for 24 hours (n = 4). (**D**) Confirmation of *Bhlhe40* overexpression in Ctrl and *B40* OE MIN6 cells by Western blotting. Data are mean ± SEM; $^{**}p < 0.01$ by unpaired two-tailed Student's *t* test. Ctrl, control.

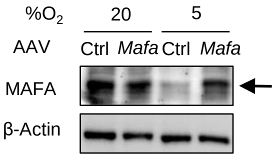


**Supplemental Figure 4 (related to Figure 4). Validation of *Mafa* overexpression mediated by AAV.** Western blot confirming *Mafa* overexpression in MIN6 cells infected with AAV-GFP (Ctrl) and AAV-*Mafa* and cultured under 20% or 5% $O_2$ for 24 hours. Ctrl, control.

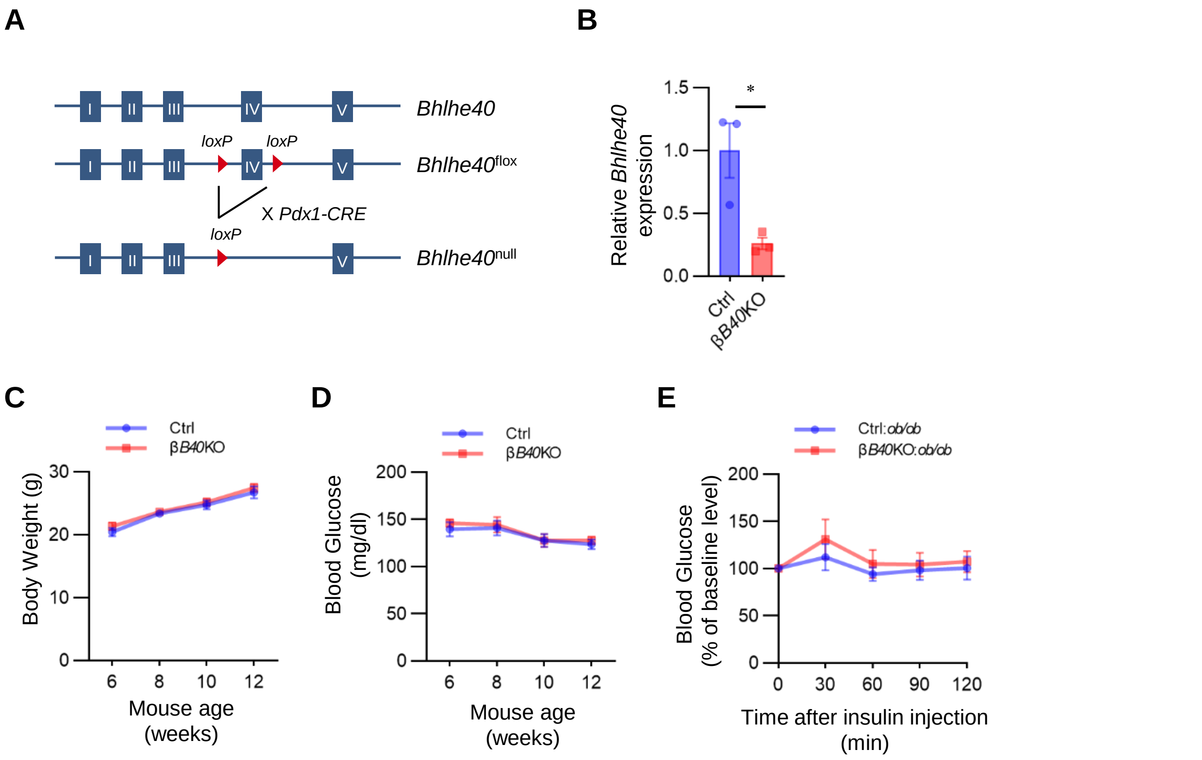


**Supplemental Figure 5 (related to Figure 6). Deficiency of BHLHE40 does not affect body weight and nonfasting blood glucose in normal mice.** (**A**) Schematic representation of conditional deletion of *Bhlhe40*. (**B**) Confirmation of BHLHE40 knockout in islets from *Bhlhe40*$^{fl/fl}$ (Ctrl) and *Pdx1*-Cre:*Bhlhe40*$^{fl/fl}$ (βB40KO) mice (n = 3) by qRT-PCR. (**C** and **D**) Body weight (**C**) and nonfasting blood glucose (**D**) in Ctrl and βB40KO mice (n = 7 and n = 9, respectively). (**E**) Insulin tolerance test of Ctrl:*ob/ob* and βB40KO:*ob/ob* mice (n = 9 and n = 8, respectively; 7 weeks old). Data are mean ± SEM; *p < 0.05 by unpaired two-tailed Student's *t* test. Ctrl, control.

**Supplemental Table 1. Oligo DNAs used in this study.**

| qPCR Primers | FORWARD | REVERSE |
|---|---|---|
| ***Actb*** | GGCCGGGACCTGACAGACTA | AGGAAGAGGATGCGGCAGTG |
| ***Bhlhe40*** | CTGCAGCCCTCTCCAGCTTC | TGGAGCAAGGCCGAAGAGTC |
| ***MafA*** | CAGCAGCGGCACATTCTG | GCCCGCCAACTTCTCGTAT |
| ***Napa*** | GCCAACAAGTGTCTGCTGAA | GTCAATGCAGAAGTGGCAGA |
| ***Neurod1*** | TCCCACGTCTTCCACGTCAA | TTTCAAACTCGGCGGATGGT |
| ***Nkx2-2*** | CTTGGTCAGGGACGGCAAAC | GGTGCTGGCCGAGCTGTACT |
| ***Nkx6-1*** | ACTTGGCAGGACCAGAGAGA | AGAGTTCGGGTCCAGAGGTT |
| ***Ppargc1a*** | GAAATCCGAGCGGAGCTGAA | GAATAGGGCTGCGTGCCATC |
| ***Slc30a8*** | AGGTGGTGGGTGGACACGTT | AAAAGGCGCTCACAGGCAAG |
| ***Stx1a*** | GAAGGTCTGAACCGCTCATC | ATTCCTCACTGGTCGTGGTC |
| ***Stxbp1*** | CTGAAGAACGGTATCACTGA | GTAGGTCTGCTCACTGATAC |
| ***Syt7*** | ACCTCAAAGCCATGGACATC | GGCTGAGCTTGTCTTTGTCC |
| ***Tbp*** | CCCCTTGTACCCTTCACCAAT | GAAGCTGCGGTACAATTCCAG |
| ***Vdr*** | GACCGCCTATCCAACACACT | ATCTCATTGCCGAACACCTC |

| ChIP-qPCR Primers | FORWARD | REVERSE |
|---|---|---|
| ***MafA* enhancer Site A** | CTCTCCCTGCCTGAGCCTTG | CACTCGATCCATGGCACCTG |
| ***MafA* enhancer Site B** | TGTGAGGGTGAGGGGCTAGG | CCTGCAGGAGAGGGGACAAA |
| ***MafA* enhancer Site C** | TGTGTCTGGCAGCCTTGCTC | AGAGGGGAAAGGGACCAGGA |
| ***MafA* enhancer Site D** | TCCCATGAGCACCAGCAGAG | TAGGCCCAATGGCACTGGAT |
| ***MafA* enhancer Site TF binding** | CACCCCAGCGAGGGCTGATTTAATT | AGCAAGCACTTCAGTGTGCTCAGTG |
| ***Tbp*** | CCCCTTGTACCCTTCACCAAT | GAAGCTGCGGTACAATTCCAG |
